# Multivalent coiled-coil interactions enable full-scale centrosome assembly and strength

**DOI:** 10.1101/2023.05.15.540834

**Authors:** Manolo U. Rios, Małgorzata A. Bagnucka, Bryan D. Ryder, Beatriz Ferreira Gomes, Nicole Familiari, Kan Yaguchi, Matthew Amato, Łukasz A. Joachimiak, Jeffrey B. Woodruff

## Abstract

During mitotic spindle assembly, microtubules generate tensile stresses on pericentriolar material (PCM), the outermost layer of centrosomes. The molecular interactions that enable PCM to assemble rapidly and resist external forces are unknown. Here we use cross-linking mass spectrometry to identify interactions underlying supramolecular assembly of SPD-5, the main PCM scaffold protein in *C. elegans*. Crosslinks map primarily to alpha helices within the phospho-regulated region (PReM), a long C-terminal coiled-coil, and a series of four N-terminal coiled-coils. PLK-1 phosphorylation of SPD-5 creates new homotypic contacts, including two between PReM and the CM2-like domain, and eliminates numerous contacts in disordered linker regions, thus favoring coiled-coil-specific interactions. Mutations within these interacting regions cause PCM assembly defects that are partly rescued by eliminating microtubule-mediated forces. Thus, PCM assembly and strength are interdependent. *In vitro*, self-assembly of SPD-5 scales with coiled-coil content, although there is a defined hierarchy of association. We propose that multivalent interactions among coiled-coil regions of SPD-5 build the PCM scaffold and contribute sufficient strength to resist microtubule-mediated forces.

## INTRODUCTION

Regulated assembly of pericentriolar material (PCM) is critical for proper mitotic cell division, and PCM dysfunction is linked to various cancers and developmental disorders such as microcephaly and primordial dwarfism (Goundiam and Basto, 2021). PCM is typically nucleated by the centrioles at centrosomes; however, PCM can also assemble independently (Garbrecht et al., 2021; Magescas et al., 2021; Schuh and Ellenberg, 2007). PCM is a micron-scale, supramolecular mass of protein that nucleates and anchors microtubule arrays (Conduit et al., 2015; Vasquez-Limeta and Loncarek, 2021; Woodruff et al., 2014). Self-assembly of large coiled-coil proteins, such as SPD-5 (*C. elegans*) and Centrosomin (*D. melanogaster*) form the underlying PCM “scaffold” (Feng et al., 2017; Fong et al., 2008; Hamill et al., 2002; Megraw et al., 1999; Woodruff et al., 2015). This scaffold recruits “client” proteins that regulate PCM assembly, material properties, and function. The PCM clients γ-tubulin, TPX2, and ch-TOG concentrate α/β tubulin and nucleate microtubules (King and Petry, 2020; Moritz et al., 1995; Roostalu et al., 2015; Wieczorek et al., 2015; Zheng et al., 1995). Polo Kinase, SPD-2/Cep192, and Aurora A Kinase potentiate PCM scaffold assembly (Conduit et al., 2010; Conduit et al., 2014; Haren et al., 2009; Woodruff et al., 2015), but they are not essential in certain developmental contexts (Garbrecht et al., 2021; Magescas et al., 2021). PP2A-B55a Phosphatase weakens PCM and promotes its disassembly at the end of each cell cycle in *C. elegans* embryos (Enos et al., 2018; Magescas et al., 2019; Mittasch et al., 2020).

During mitotic spindle assembly and positioning, the PCM must bear microtubule-dependent forces that create tensile stresses, yet the molecular basis is unclear. Motor proteins anchored at the plasma membrane walk along astral microtubules to generate cortically-directed forces (Laan et al., 2012; Pecreaux et al., 2006). Spindle-microtubule-localized motor proteins also pull at PCM (Dumont and Mitchison, 2009). During anaphase in *C. elegans* embryos, these forces fracture and disperse the PCM scaffold and accelerate its disassembly (Enos et al., 2018; Magescas et al., 2019). Non-invasive nano-rheology demonstrated that PCM strength and ductility decline sharply during the metaphase-to-anaphase transition (Mittasch et al., 2020), indicating that PCM material properties are tuned during the cell cycle. These experiments also revealed that PCM is more susceptible to shear stress when PLK-1 activity is inhibited, suggesting that phosphorylation strengthens the PCM scaffold (Mittasch et al., 2020). However, PLK-1 inhibition does not promote premature PCM fracture, making unclear if phosphorylation is critical for PCM resistance to native tensile stresses. Therefore, to reveal the molecular basis of PCM force resistance, it is important to identify molecular connections that prevent PCM fracture under native microtubule-mediated forces.

Reconstitution experiments revealed that full-length SPD-5 and Centrosomin are each sufficient to multimerize into micron-scale assemblies (Feng et al., 2017; Woodruff et al., 2017; Woodruff et al., 2015). Analysis of SPD-5 fragments showed an interaction between the CM2-like domain a central region containing coiled-coil domains and PLK-1 phosphorylation sites (PReM region)(Nakajo et al., 2022). Similarly, an interaction between two coiled-coil domains in Centrosomin (LZ and CM2) is critical for PCM assembly in flies (Feng et al., 2017). While these domains are clearly important for PCM assembly, there are likely additional interactions needed to make a micron-scale scaffold. Li et al. (2012) demonstrated that a minimum valence of 3 (i.e., 3 distinct binding domains) is required to achieve 3-dimensional protein phases and network structures. Increasing the valence beyond this value further decreases the threshold concentration for network formation. Phase separation of a synthetic peptide at physiologically possible concentrations (<50 μM) required a valence of 5. The molecular interactions sufficient for micron-scale PCM scaffold assembly have not been comprehensively characterized in any species. Nor is it known if homotypic interactions between scaffold proteins are important for scaffold assembly, force resistance, or both.

Here, we map the molecular interactions underlying assembly of the SPD-5 scaffold using cross-linking mass spectrometry (XL-MS) and test their importance for PCM assembly and force resistance in *C. elegans* embryos. Our results suggest that SPD-5 assembles largely through interactions mediated by coiled-coil domains distributed throughout the protein. PLK-1 phosphorylation of SPD-5 eliminates many disordered linker-derived contacts and creates few new contacts, including one between the CM2-like and PReM regions. Mutating mapped interacting regions in the N-terminus (a.a. 1-566) and a long coiled-coil (a.a. 734-918) of SPD-5 impair PCM assembly *in vivo* and make PCM weak and susceptible to microtubule-dependent pulling forces. In a companion paper, we highlight the role of a structured coiled-coil domain in SPD-5 (a.a. 541-677) that is critical for PCM assembly and strength (Rios et al., 2023). Our results reveal that PCM assembly and material properties are intertwined: PCM must be able to resist microtubule-dependent forces to assemble properly. We conclude that multivalent coiled-coil interactions between SPD-5 proteins enable both full-scale PCM assembly and force resistance.

## RESULTS

### XL-MS reveals interactions during SPD-5 self-assembly

SPD-5 is essential for PCM assembly in *C. elegans* embryos (Hamill et al., 2002) and sufficient to form micron-scale assemblies that recruit client proteins and nucleate microtubule asters *in vitro* (Woodruff et al., 2017; Woodruff et al., 2015)(Figure 1A). Alphafold predicts that SPD-5 contains 14 alpha helices connected by disordered linker regions (Jumper et al., 2021) (Fig. 1B). MARCOIL (50% threshold; MTK matrix) predicts that 9 disconnected regions within the alpha helices form coiled-coils. PLK-1 phosphorylation accelerates SPD-5 multimerization *in vitro* and is required for full-scale PCM assembly in *C. elegans* embryos (Woodruff et al., 2015). To reveal how phosphorylation regulates SPD-5 assembly, and to identify the most physiologically relevant interaction sites, we performed XL-MS on *in vitro* SPD-5 scaffolds assembled in the presence of constitutively-active (CA) or kinase-dead (KD) PLK-1. We used the zero-length cross-linker DMTMM, which captures interactions between primary amines of lysines and the carboxylates of aspartic or glutamic acids. For this study, we report only interactions between SPD-5 molecules in the multimeric state.

**Figure 1.**
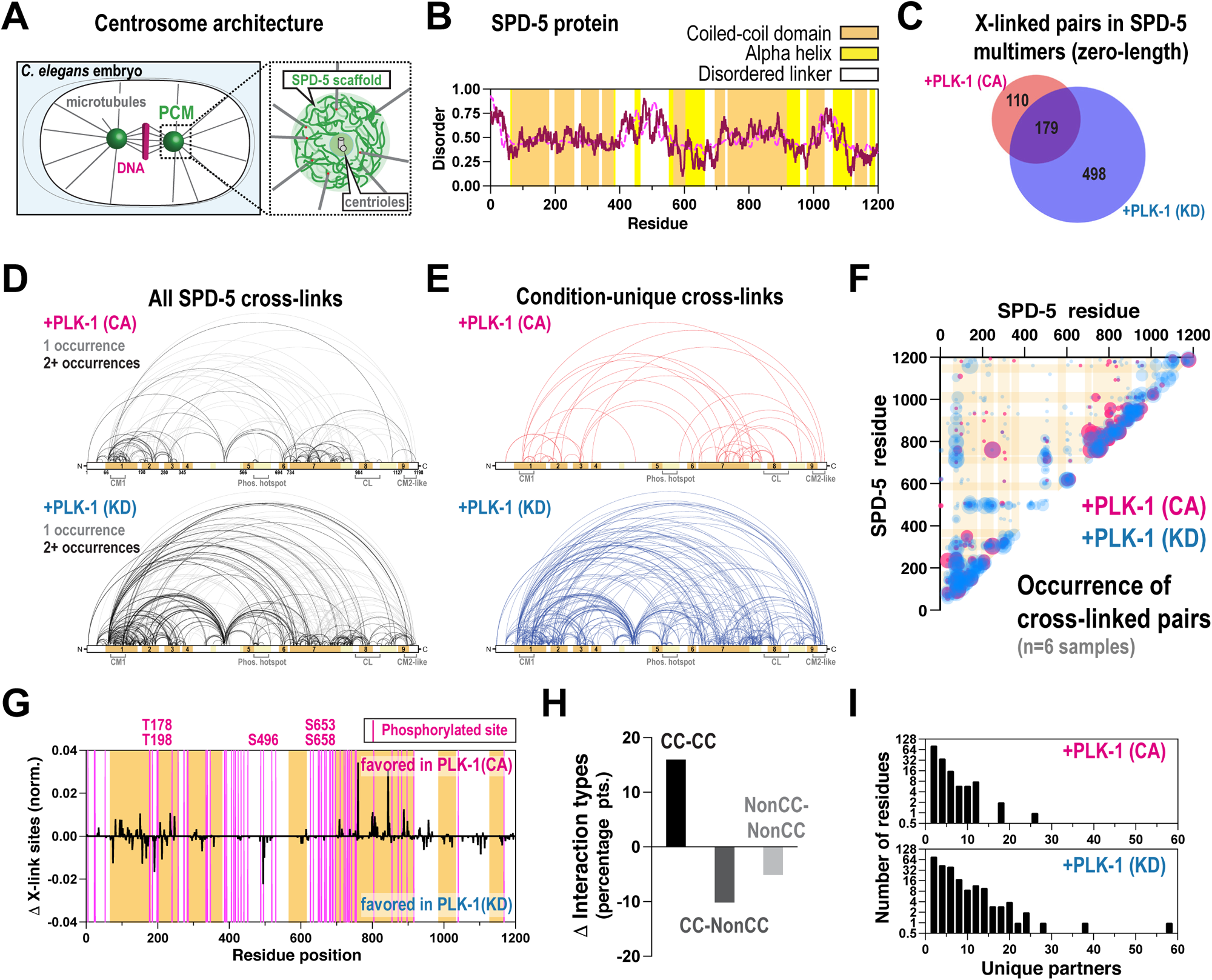
PLK-1 phosphorylation alters SPD-5 homotypic interactions. A. Centrosome architecture in a *C. elegans* embryo (top panels). B. Secondary structural analysis of SPD-5 featuring predicted coiled-coil domains (MARCOIL 50%), alpha helices (Alphafold, >70% confidence) and disorder scores (IUPRED2 score in purple and ANCHOR score in magenta). C. Purified SPD-5 was dephosphorylated, then assembled in the presence of constitutively active (CA) or kinase-dead (KD) PLK-1. Samples were cross-linked using DMTMM, then SPD-5 multimers were analyzed by mass spectrometry (n= 6 replicates). The venn diagram shows the number of non-redundant cross-linked pairs found in each condition corresponding to SPD-5 homotypic interactions. D. Map of DMTMM-induced crosslinks found per condition, pooled from 6 replicates (+PLK-1(CA), n= 289 crosslinks; +PLK-1(KD) n = 677 crosslinks). Crosslinks appearing in 2 or more samples are shown with black lines. SPD-5 domain legend: CM1, g-tubulin binding domain; Phos. Hotspot, region containing essential PLK-1 sites; CL, centriole-localization domain; CM2-like, essential region for SPD-5 assembly. E. Crosslinks found only in CA or KD samples. F. Quantification and location of cross-linked pairs in the pooled CA (red) or KD (blue) samples. Bubble size indicates number of replicates containing a given cross-linked pair. G. The percentage of cross-linked pairs involving predicted coiled-coil domains (CC) or linker domains was determined for each condition. Shown is the difference in percentage points for each category between the CA and KD samples. See Figure S1 original data. H. The total number of times a residue was identified in a cross-linked pair was calculated for each condition. Data were normalized to account for overall differences in identified crosslinks. Shown is the difference between CA and KD samples. See Figure S1 for original data. I. For each residue identified in a cross-linked pair, the number of unique partners was calculated.

Analysis of 6 replicates per condition identified 2.3-fold less cross-linked pairs in the CA vs. KD samples (289 vs. 677 non-duplicate pairs)(Figure 1C,D; see Supplemental Data Set 1 for a list of all crosslinks). This result implies that phosphorylation reduces SPD-5 homotypic interactions overall. In both conditions, interactions were concentrated in numerous coiled-coil domains and not the linkers (Figure 1D,F and S1A). We identified pairs unique to CA samples, indicating that specific interactions are gained via phosphorylation. Other pairs were unique to the KD samples, indicating that these interactions are lost when SPD-5 is phosphorylated (Figure 1E). Interactions unique to the phosphorylated state were spatially close to those found in the unphosphorylated state: 31% of CA-unique crosslinks were within 5 a.a. of KD-unique crosslinks (Figure 1F). The CA sample contained two unique crosslinks (D349-K1183 and E665-K1160) between the PReM-containing (a.a. 272-732) and CM2-like (a.a. 1061-1198) regions. This result is consistent with pull-down experiments showing that PLK-1 phosphorylation enhances the binding between reconstituted PReM and CM2 regions (Nakajo et al., 2022).

We next assessed how phosphorylation affects motif preference during SPD-5 scaffold assembly. We used MS to identify SPD-5 residues phosphorylated by PLK-1 (Figure 1G for a map, Supplemental Data Set 2 for a list). After normalizing for differences in overall cross-link number, we found that phosphorylation of SPD-5 increases its preference for coiled-coil interactions and reduces the number of linker interactions (Figure 1H and S1A,B). The most prominent gains were detected in coiled-coils 1, 2 and 7. The most prominent losses were at residues K191 and K495. The change at residue K191 is noteworthy as it is close to PLK-1 phosphorylation sites (T178 and T198) that regulate docking of the γ-tubulin complex (Ohta et al., 2021). Interacting residues had overall fewer observed crosslinked partners in the phosphorylated state (Figure 1I and S1C). Distributions of sequence separation between cross-linked sites were similar between samples, suggesting that phosphorylation does not induce changes in the register between the coiled-coils (Figure S1D). We conclude that PLK-1 phosphorylation regulates SPD-5 assembly by tweaking its interaction landscape in three ways: 1) promoting interactions between the PReM and CM2 domains, 2) blocking disordered linker interactions, and 3) shifting the location of coiled-coil contacts to proximal sites. Our results are consistent with the model wherein PLK-1 phosphorylation of SPD-5 eliminates weaker, less-specific linker contacts to favor more specific and stronger interactions between the coiled-coil domains. These data are consistent with the “relief of auto-inhibition” models proposed for how Polo Kinase potentiates SPD-5 binding to the γ-tubulin complex and Centrosomin assembly and binding to the γ-tubulin complex (Feng et al., 2017; Ohta et al., 2021; Tovey et al., 2021).

Our analyses demonstrate that crosslinks in the SPD-5 multimers primarily occur between coiled-coil domains and not linkers, suggesting that coiled-coil domains are the primary drivers of scaffold assembly. We mapped interactions within previously characterized CM2-like and PReM domains, supporting the validity of our approach. Our crosslinks are also sufficient to reveal new structural motifs, including an alpha helical hairpin within the PReM domain (see our companion paper; Rios et al., 2023). In addition, we identified previously uncharacterized interaction sites, including two hotspots: a C-terminal, long coiled-coil domain (CC-Long; a.a. 734-918) and four coiled-coil domains in the N-terminus. While XL-MS allows identification of contact sites, it cannot indicate whether these contact sites are required for protein assembly and function. In a companion study, we demonstrate that the helical hairpin in SPD-5 is essential for PCM assembly and strength *in vivo* (Rios et al., 2023). We found that SPD-5 mutants lacking the CM2-like domain (a.a. 1061-1198) express poorly in embryos, consistent with prior observations (Nakajo et al., 2022), making it difficult to evaluate the necessity of this domain. Therefore, in this study, we focused on testing the importance of the newly identified coiled-coil interaction motifs for PCM assembly and strength in *C. elegans* embryos.

### A long, C-terminal coiled-coil domain is essential for SPD-5 assembly and strength

Alphafold predicts that SPD-5 contains a long alpha helix spanning a.a. 739-957 (confidence score >70); MARCOIL (50% threshold) predicts that a.a. 734 -918 constitutes a coiled-coil domain, which we term CC-Long. Our XL-MS identified 200 cross-linked pairs where at least one residue resided within this region. To test the importance of CC-Long for PCM assembly and strength, we used CRISPR to delete a.a. 734-918 in a strain expressing wild-type SPD-5 endogenously tagged with RFP (RFP::SPD-5(FL)). Embryos expressing RFP::SPD-5(FL) built PCM that was spherical, as expected (Figure 2A-D). Embryos expressing RFP::SPD-5(Δ734-918) were viable and could build PCM that directed bipolar spindle formation (Figure S2A), but PCM was smaller and more irregular in shape compared with the control (Figure 2A-D). PCM ruptured and disassembled in late anaphase in control embryos, whereas PCM fragmented prematurely in *spd-5(*Δ*734-918)* embryos (Figure 2B,E). These phenotypes were not due to different expression of the two constructs, as levels of both wild-type and mutant proteins were equivalent (Figure S2B). To test if the irregular PCM morphology was due to an inability to resist native microtubule-mediated pulling forces, we eliminated forces by adding nocodazole to embryos. Nocodazole treatment increased the mean intensity of PCM-localized RFP::SPD-5(Δ734-918) and restored the spherical shape of the PCM scaffold in metaphase (Figure 2A-C). These data indicate that irregular PCM morphology is caused by microtubule-mediated pulling forces, suggesting that loss of CC-Long leads to material weakness in the PCM scaffold. While nocodazole treatment rescued PCM circularity, it only increased PCM assembly ∼1.4-fold compared to the DMSO control, which was well below wild-type levels (Figure 2D). Thus, CC-Long is also required for full-scale SPD-5 assembly even in the absence of pulling forces.

**Figure 2.**
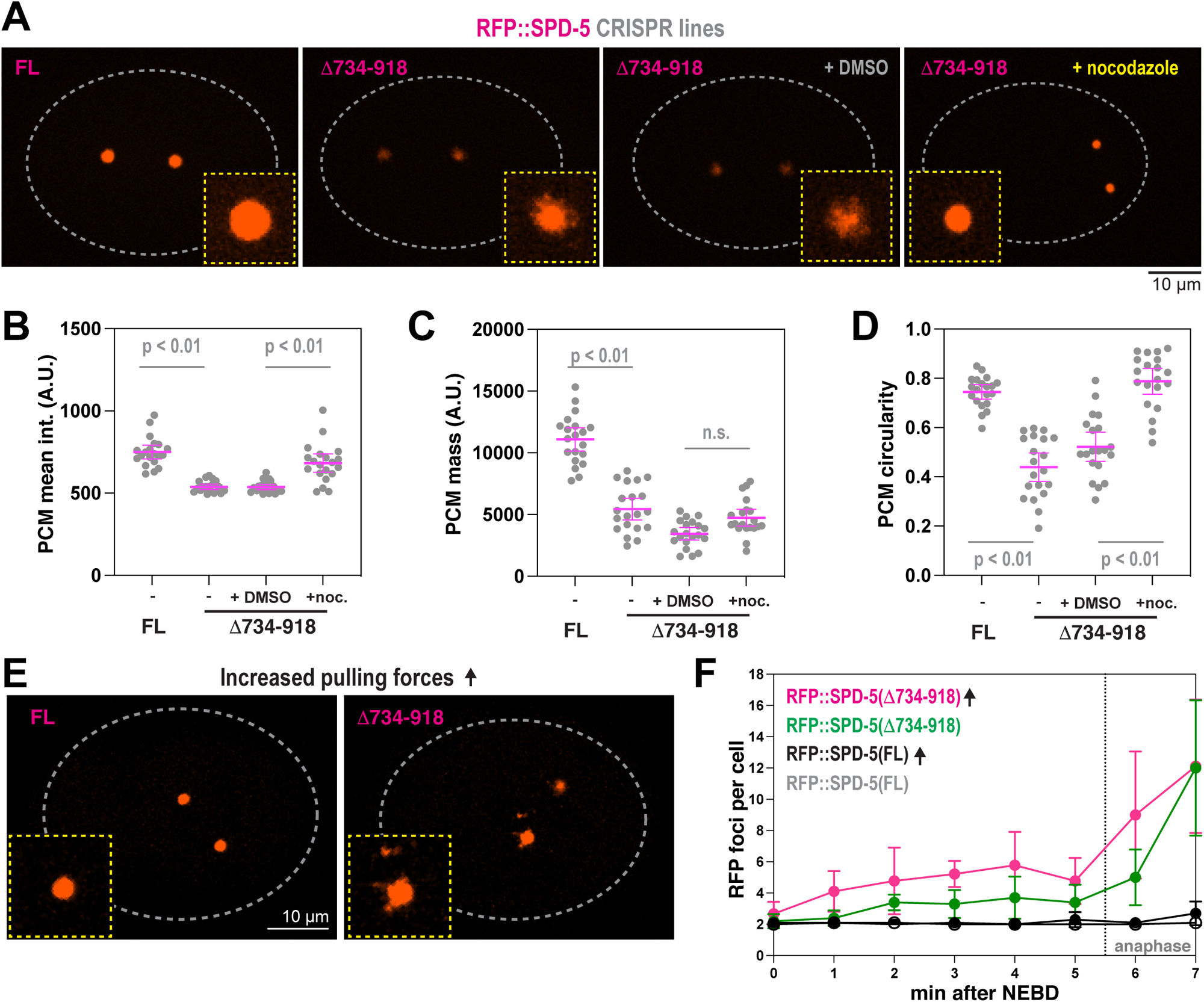
A long central coiled-coil motif in SPD-5 is essential for PCM strength and assembly. A. Endogenous RFP-tagged SPD-5 was modified by CRISPR to delete residues 734-918. Embryos expressing the unmodified (WT) and mutant RFP::SPD-5 were visualized with fluorescence confocal microscopy. 20 μM nocodazole or 1% DMSO were introduced via laser puncture of the eggshell. B. Quantification of mean intensity of RFP signal localized at PCM during metaphase. Mean +/- 95% C.I.; N = 20 centrosomes; p value from a Kruskal-Wallis test. C. Quantification of fluorescence integrated density of RFP signal localized at PCM during metaphase. Mean +/- 95% C.I.; N = 20 centrosomes; p value from a Kruskal-Wallis test. D. Quantification of PCM circularity during metaphase in indicated strains. Mean +/- 95% C.I.; N = 20 centrosomes; p value from a Kruskal-Wallis test. E. Embryos were treated with RNAi against *csnk-1*, which increases microtubule-mediated pulling forces. Insets show zoomed in images of centrosomes. F. PCM fragmentation from nuclear envelope breakdown onward in one-cell embryos. Mean +/- 95% C.I.; N = 10 embryos. Arrows indicate embryos were treated with *csnk-1* RNAi.

Having shown that ablating pulling forces rescues PCM circularity and assembly defects in *spd-5(*Δ*734-918)* embryos, we next asked whether increasing pulling forces would worsen the phenotype. RNAi knockdown of casein kinase 1 gamma (CSNK-1), which increases cortical pulling forces by ∼1.5-fold (Panbianco et al., 2008), exacerbated the PCM fragmentation phenotype in embryos expressing SPD-5(Δ734-918) but not SPD-5(FL) (Figure 2E,F). Taken together, these results indicate that PCM built with SPD-5(Δ734-918) is too weak to stay connected under physiological pulling forces. We conclude that that CC-Long is required for the SPD-5 scaffold to self-assemble and generate enough strength to resist microtubule-mediate pulling forces.

### The N-terminus of SPD-5 is essential for full-scale PCM assembly and strength

The N-terminus of SPD-5 (a.a. 1-566) contains four predicted coiled-coil domains, the first of which contains a CM1-like region that recruits the γ−tubulin complex (Ohta et al., 2021). Our XL-MS identified 256 cross-linked pairs where at least one residue resided within a.a. 1-566, raising the possibility that this region could mediate SPD-5 scaffold assembly and strength. We used MoSCI to create a transgenic version of SPD-5 missing the N-terminus (GFP::SPD-5(566-1198)) and compared it with transgenic full-length SPD-5 (GFP::SPD-5(FL)). These transgenes are resistant to RNAi knockdown of endogenous *spd-5* (Figure 3A and S2C). After RNAi depletion of endogenous *spd-5*, GFP::SPD-5(566-1198) was sufficient to build PCM that was, on average, 17-fold less massive than in control embryos (Figure 3B,C). The mutant embryos did not develop to the larval stage under these conditions (100% lethality; Figure S2D). We conclude that the N-terminus of SPD-5 is essential for full-scale PCM assembly.

**Figure 3.**
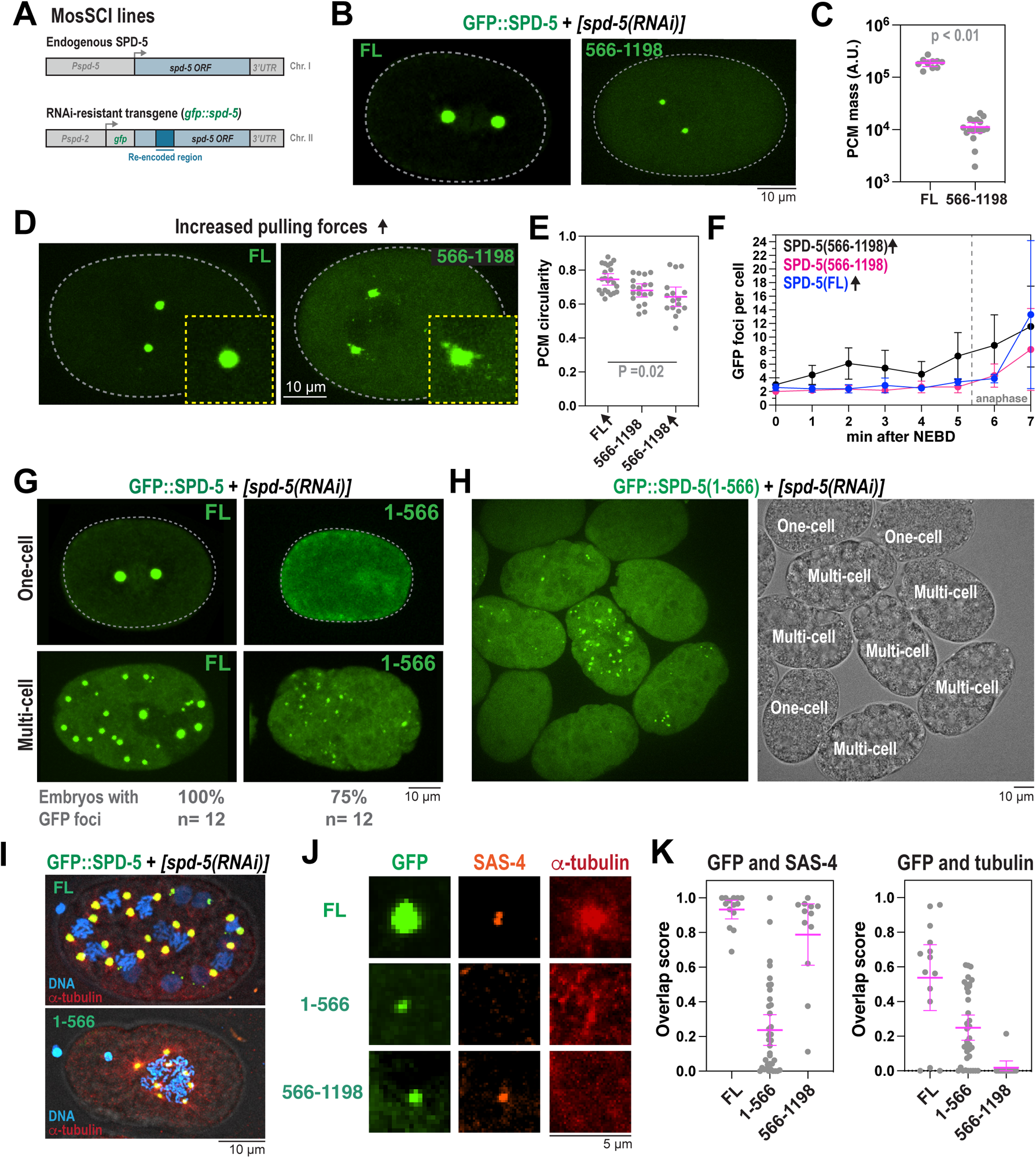
The coiled-coil-containing N-terminus of SPD-5 is required for PCM strength and assembly. A. MosSCI-generated transgenes are expressed from chromosome II and are re-encoded to be resistant to RNAi knockdown. Endogenous *spd-5* is expressed from chromosome I. B. Fluorescence confocal images of embryos expressing full-length (a.a. 1-1198; FL) or truncated (a.a. 566-1198) *spd-5* transgenes after knockdown of endogenous SPD-5 by RNAi. C. Quantification of fluorescence integrated density of GFP signal localized at PCM during metaphase. Mean +/- 95% C.I.; N = 10 (FL) and 17 (566-1198) centrosomes; p value from a Mann-Whitney test. D. Images of *csnk-1(RNAi)* embryos expressing endogenous *spd-5* and transgenic *gfp::spd-5*. E. Quantification of PCM circularity during metaphase in (B,D). Arrows indicate embryos were treated with *csnk-1* RNAi. Mean +/- 95% C.I.; N = 16-22 centrosomes; p value from a Kruskal-Wallis test. F. PCM fragmentation from nuclear envelope breakdown onward in one-cell embryos. Mean +/- 95% C.I.; N = 6-10 embryos. Arrows indicate embryos treated with *csnk-1* RNAi. G. Images of one-cell and multi-cell embryos expressing full-length (FL) or truncated SPD-5 (a.a. 1-566). The number of multi-cell embryos displaying GFP-positive foci is indicated below. H. Embryos expressing GFP::SPD-5(1-566) in the absence of endogenous SPD-5. Fluorescence image on the left, DIC image on the right. Cells are labeled based on developmental stage. I. Immunofluorescence of transgenic embryos stained for alpha-tubulin and DNA. Embryos were lightly fixed to preserve the native GFP fluorescence. J. Immunofluorescence of transgenic embryos stained for a centriolar marker (SAS-4) and alpha tubulin. K. (Left) Overlap scores between anti-SAS-4 and GFP::SPD-5 signal. (Right) Overlap scores between anti-alpha-tubulin and GFP::SPD-5 signal. Mean +/- 95% C.I.; n = 12-36 GFP-positive foci.

Treatment with nocodazole had no effect on PCM assembly in embryos expressing GFP::SPD-5(566-1198). This result is expected, since the N-terminus of SPD-5 anchors microtubules though the γ-tubulin complex (Ohta et al., 2021); thus, pulling forces are not transmitted and are not expected to strain the PCM. We reasoned that if SPD-5(566-1198) is defective in generating strength, it might disrupt the integrity of PCM built with endogenous SPD-5. In embryos expressing endogenous SPD-5, GFP::SPD-5(566-1198) localized to PCM but did not cause PCM fragmentation or premature disassembly (Figure S2H). To test whether increasing pulling forces would reveal a phenotype in this mixed scenario, we depleted CSNK-1 using RNAi. PCM prematurely ruptured and became less spherical in *csnk-1(RNAi)* embryos expressing GFP::SPD-5(566-1198), while no effect was observed for control embryos expressing GFP::SPD-5(FL) (Figure 3D-F). We conclude that SPD-5(566-1198) interferes with proper assembly of endogenous SPD-5, resulting in structurally weak PCM. Our results suggest that interactions mediated by SPD-5’s N-terminal helical domains are essential to achieve full-scale PCM assembly and strength.

Our finding that the N-terminus of SPD-5 is necessary for full-scale PCM assembly is seemingly inconsistent with a prior study showing that a N-terminal SPD-5 fragment does not assemble in one-cell embryos (Nakajo et al., 2022). To investigate this matter, we used MosSCI to create embryos expressing N-terminal SPD-5 (a.a. 1-566) tagged with GFP. In the absence of endogenous SPD-5, GFP::SPD-5(1-566) was not sufficient to form PCM in one-cell embryos or rescue viability, consistent with the prior study (Figure 3G and S2D). However, in further developed multi-cell embryos, GFP::SPD-5(1-566) assembled into foci detectable by confocal microscopy (Figure 3G). This phenomenon showed cell-to-cell variability and was most prominent in embryos depleted of endogenous SPD-5 (Figure 3H). We noticed that transgene expression increased after knockdown of endogenous SPD-5 and during development (Figure S2E-G). Thus, we speculate that SPD-5(1-566) assembly *in vivo* is highly sensitive to protein concentration and only occurs after crossing a critical threshold, consistent with polymer percolation theory (Harmon et al., 2017; Li et al., 2012).

Immunofluorescence revealed that GFP::SPD-5(1-566) foci could nucleate microtubules and did not localize with the centriolar marker SAS-4, indicating that these structures represent semi-functional, PCM-like assemblies but not true centrosomes (Figure 3I-K). Endogenous SPD-5 was not detected at GFP::SPD-5(1-566) foci, indicating efficient RNAi knockdown (Figure S2H). Conversely, GFP::SPD-5(566-1198) foci co-localized with SAS-4 and not microtubules (Figure 3J,K). This is consistent with previous studies showing that the N-terminus of SPD-5 binds γ-tubulin complexes (Ohta et al., 2021), while the C-terminus is required for centriole binding (Nakajo et al., 2022). These results suggest that N-terminal coiled-coils mediate both self-assembly and the microtubule-nucleation capacity of SPD-5. We conclude that assembly of full-sized, functional PCM scaffold requires multiple domains in both the N- and C-termini of SPD-5.

### SPD-5 scaffold assembly scales with coiled-coil content

Our crosslinking data and mutational analyses, as well as previous studies, indicate that SPD-5 contains several coiled-coil-rich modules that mediate self-assembly. This type of multivalent architecture is common among proteins that form supramolecular networks, gels, and liquid droplets (Banani et al., 2017; Li et al., 2012). If SPD-5 assembles through multivalent interactions, then multiple SPD-5 domains should be sufficient to self-assemble or co-assemble with each other. We therefore purified 9 separate fragments of SPD-5 and assayed multimerization *in vitro* (Figure 4A and S3A). Full-length SPD-5 (FL) and fragments that were predicted to contain coiled-coil domains showed strong alpha-helical signatures using circular dichroism spectroscopy (Figure 4B). When diluted into physiological salt (150 mM KCl) and a macromolecular crowding agent (7.5% PEG, 3,000 MW), pure SPD-5(FL) spontaneously assembles into micron-scale, spherical condensates (Woodruff et al., 2017). All fragments except F23, which lacks coiled-coil domains, assembled into supramolecular structures detectable by light microscopy; yet, none could assemble to the same extent as FL (Figure 4C). Only FL, F24 (a.a. 566-1198), F26 (a.a. 730-1198), and F28 (CC-long; Δ734-918) proteins were sufficient to assemble spherical, micron-scale condensates. Other fragments instead assembled into smaller, irregular aggregates. At equimolar concentrations, SPD-5 assembly scaled non-linearly with the molecular mass of coiled-coil domains (R^2^=0.78; Figure 4D). These data indicate that no fragment can fully recapitulate the assembly properties of full-length SPD-5. Thus, a high degree of valency is required to achieve proper scaffold mass and morphology. Our results also indicate that C-terminal fragments assemble better than N-terminal fragments (e.g., compare F24 and F21), which mirrors our findings in one-cell embryos (Figure 3). We saw a striking reduction in assembly of F26 vs. F24, indicating the importance of a.a. 566-730 for SPD-5 multimerization. This result is consistent with our companion study showing that this region comprises an alpha helical hairpin that is essential for PCM assembly and strength (Rios et al., 2023). We conclude that coiled-coil domains drive SPD-5 multimerization, but the domains do not play equivalent roles.

**Figure 4.**
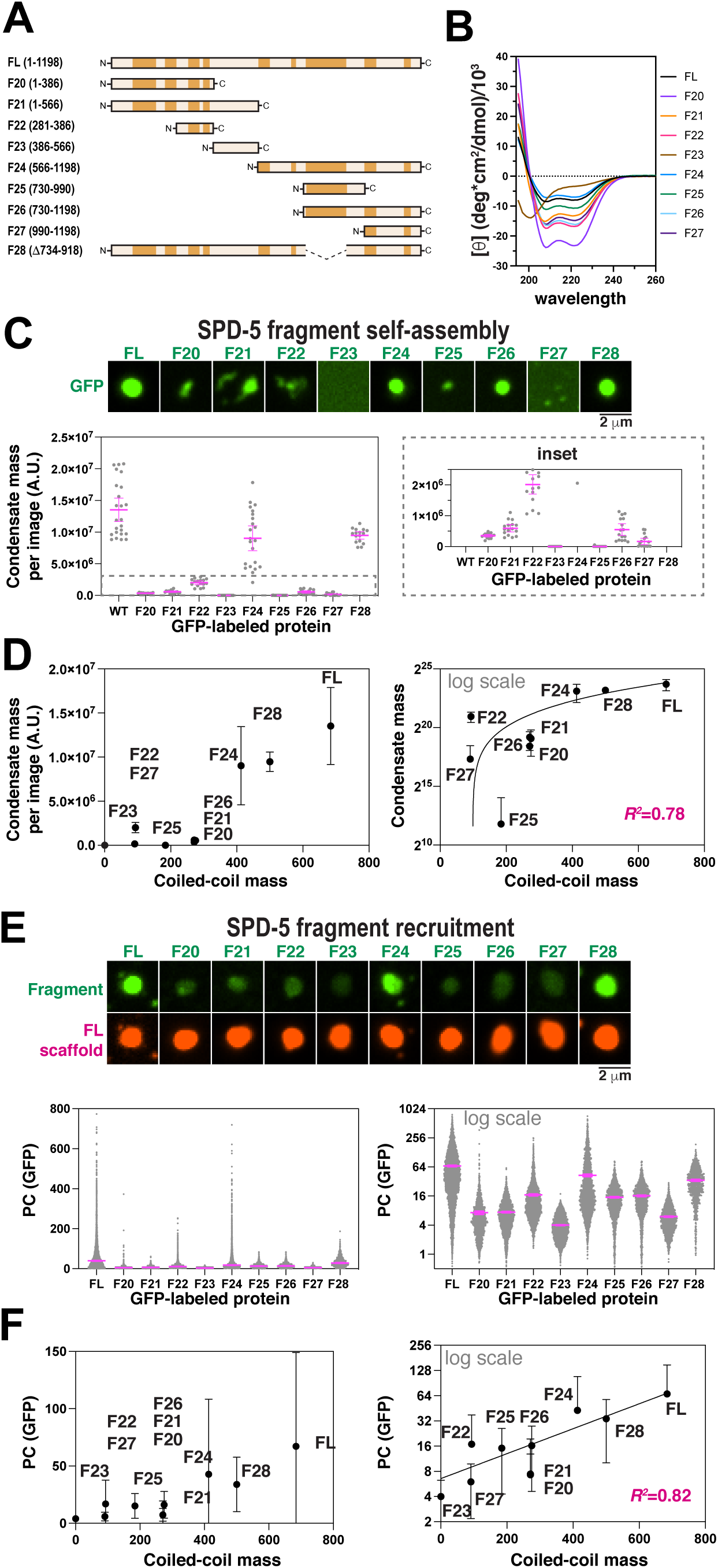
Multiple SPD-5 regions are sufficient to form micron-scale assemblies *in vitro*. A. Domain maps of 9 different SPD-5 variants. B. Circular dichroism spectroscopy of purified SPD-5 fragments. C. SPD-5 self-assembly assay. 500 nM of purified GFP-labeled SPD-5 proteins were incubated for 10 min in 7.5% (w/v) PEG-3350 then imaged. Mean +/- 95% C.I.; n = 11-22 images per condition D. Comparison of *in vitro* condensate mass with coiled-coil content. Mean +/- 95% C.I.; n = 11-22 images per condition. The data are fit with a quadratic model (R^2^= 0.78). E. SPD-5 recruitment assay. 1 μM SPD-5::RFP was incubated in 7.5% PEG-3350 for two min to form condensates. 10 nM SPD-5::GFP proteins were added, incubated for 8 min, then imaged. Mean +/- 95% C.I.; n = 875-4702 condensates per condition. F. Comparison of *in vitro* recruitment with coiled-coil content. Mean +/- 95% C.I.; n = 875-4702 condensates per condition. The data are fit with an exponential model (R^2^= 0.82).

Next, we investigated which domains are sufficient for recruitment to a pre-assembled SPD-5 scaffold. We assembled condensates containing 1 μM full-length SPD-5::RFP, then added 10 nM GFP-labeled SPD-5 variants (Figure 4E). After 10 min, we analyzed the partition coefficient (*P_c_*) of the GFP signal within the RFP-labeled condensates. SPD-5(FL)::GFP partitioned strongly into existing condensates (*P_c_* = 67), as expected. All other fragments showed poor to intermediate partitioning (*P_c_* = 4-42). Like the assembly assay, we saw that C-terminal fragments were recruited better than N-terminal fragments (e.g., compare F24 and F21). We saw a 2.6-fold reduction in partitioning of F26 vs. F24, supporting the importance of a.a. 566-730 for SPD-5 self-association. Overall, there was a strong, non-linear relationship between coiled-coil content and *P_c_* (R^2^=0.82; Figure 4F). Taken together, these results demonstrate that SPD-5 assembly and recruitment scale with coiled-coil content, suggesting that coiled-coil domains are the primary drivers of SPD-5 scaffold expansion.

To investigate whether the serial linkage of coiled-coil motifs confers an advantage, we tested if increasing the concentration of a smaller fragment would make it equivalent to full-length protein in terms of assembly capacity. We thus performed SPD-5 assembly and recruitment assays using equivalent molar concentrations of coiled-coil motifs. Our results show that SPD-5(FL) outperforms the fragments (Figure S3B,C). We conclude that serial linkage of coiled-coil domains synergistically improves self-association, which could be explained as an increase in functional affinity through avidity.

## DISCUSSION

In this study, we identified molecular interactions and material design principles that enable the PCM scaffold to resist microtubule-mediated tensile stresses without fracture. Using cross-linking mass spectrometry (XL-MS), we generated a map of interactions underlying supramolecular assembly of SPD-5, the sole identified PCM scaffold protein in *C. elegans*. Interaction sites were distributed across SPD-5, with most contacts between predicted coiled-coil domains and not disordered linker regions. PLK-1 phosphorylation of SPD-5 alters its homotypic interactions by creating new interaction sites, including a link between the CM2-like and PReM domain, and by eliminating many promiscuous contacts involving disordered linker regions. In addition, we identified new interaction hot spots, including N-terminal coiled-coils and a C-terminal coiled-coil (CC-Long; a.a. 734-918). SPD-5 fragments containing interaction regions identified by XL-MS were sufficient to self-assemble into supramolecular structures *in vitro* and *in vivo*. Deletion of each region also impaired PCM assembly *in vivo*. Thus, multiple modules within SPD-5 are necessary and sufficient for PCM scaffold assembly. Expression of these mutants often led to irregular, non-spherical PCM suggestive of premature rupture and structural weakness. This phenotype was rescued by eliminating microtubule-mediated forces and exacerbated by increasing forces. Thus, PCM scaffold assembly and mechanical properties are interdependent: PCM must be strong enough to resist microtubule-mediated pulling or else it will fracture and disassemble. Our results suggest that these features emerge from the collective inter-molecular interactions between coiled-coil domains of SPD-5. Together, our *in vivo* and *in vitro* data support a model whereby multivalent coiled-coil interactions enable SPD-5 to form a micron-scale scaffold strong enough to withstand microtubule-mediated forces (Figure 5).

**Figure 5.**
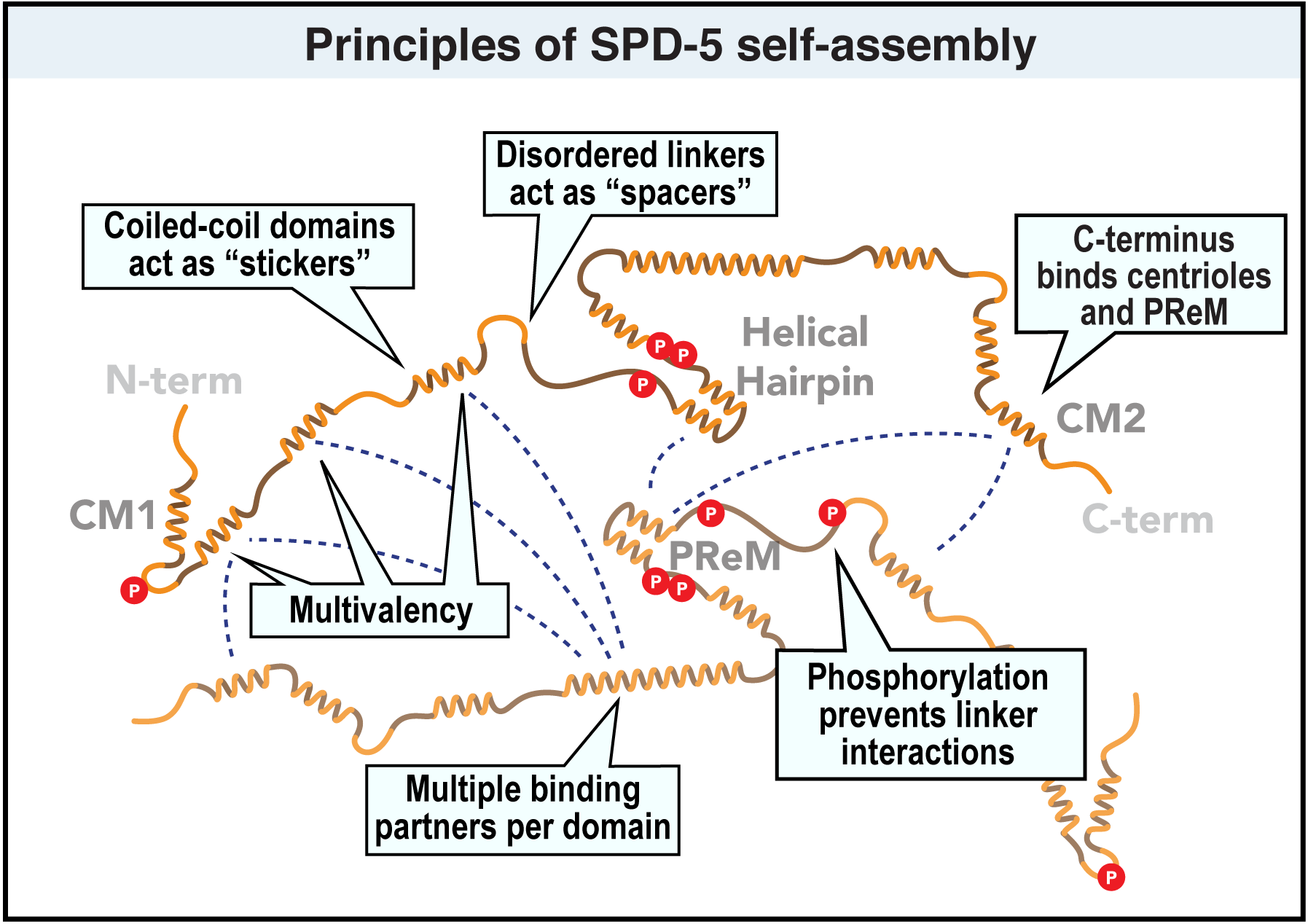
Model for SPD-5 scaffold assembly via multivalent coiled-coil interactions. SPD-5 is a largely disordered protein with coiled-coil domains separated by disordered linkers. Multiple coiled-coil domains interact to form inter-molecular connections that drive multimerization of SPD-5 into micron-scale assemblies. The linkers allow chain flexibility so that SPD-5 can sample multiple configurations. Phosphorylation of SPD-5 limits non-specific interactions between linker regions, thus favoring interactions between coiled-coil domains. The C-terminus contains a CM2-like region and centriole-localization domain. The N-terminus contains a CM1 domain that binds to gamma tubulin complexes. Strong interactions connect alpha helical hairpins to form the structural core of SPD-5 (see Rios et al., 2023). The cumulative action of these domains enables full-scale PCM assembly and strength in *C. elegans* embryos.

PLK-1 phosphorylation of SPD-5 is essential for PCM assembly, but the molecular mechanism is poorly understood. Nakajo *et al.(*2022) showed that PLK-1 phosphorylation improved the binding between SPD-5 fragments containing the PReM (a.a. 272-732) and CM2-like domains (a.a. 1061-1198). Similarly, *in vitro* reconstitution with Centrosomin, the primary PCM scaffold in flies, showed that Polo Kinase potentiates co-assembly of PReM and CM2 fragments into micron-scale networks (Feng et al., 2017). This study proposed that phosphorylation relieves auto-inhibitory interactions within Centrosomin to reveal contact sites that mediate homo-multimerization. Similar to these findings, our XL-MS analyses identified two phosphorylation-induced interactions between PReM-containing (a.a. 272-732) and CM2-like (a.a. 1061-1198) regions of SPD-5. However, the most prominent effect of PLK-1 phosphorylation was loss of interactions between disordered linker regions within SPD-5. This is especially apparent in the N-terminus, which binds to γ-tubulin complexes in a phosphorylation-dependent manner (Ohta et al., 2021). These linker-derived contacts may represent less-specific, inhibitory interactions that prevent scaffold assembly and docking of the gamma tubulin complex, consistent with the model proposed by Ohta et al. (2021). We propose that PLK-1 phosphorylation potentiates SPD-5 assembly by both eliminating auto-inhibitory linker contacts and creating new coiled-coil contacts.

Our work reveals that PCM material properties are critical for centrosome function and early embryogenesis. PCM nucleates hundreds to thousands of microtubules, which can connect to spindle-, chromatin-, and cortex-localized motors that generate force (Dumont and Mitchison, 2009; Redemann et al., 2017). These forces underlie PCM flaring seen in *D. melanogaster* and help rupture and disassemble PCM during anaphase in *C. elegans* (Enos et al., 2018; Magescas et al., 2019; Megraw et al., 2002). Such flaring or rupture is never observed during mitosis prior to late anaphase likely because PCM is materially strong at this time (Mittasch et al., 2020). The molecular interactions that enable PCM strength are not fully understood. Laser ablation of centrioles causes PCM fragmentation in a microtubule-dependent manner in *C. elegans* embryos (Cabral et al., 2019), suggesting that connection to centrioles is a major determinant of PCM force resistance. Our current study and a companion study (see Rios et al., 2023) identify key modules within SPD-5 that enable PCM strength on the mesoscale. Embryos expressing SPD-5 mutatnts lacking CC-long (a.a. 734-918)(this study) or the central hairpin (a.a. 610-640)(see Rios et al., 2023) display irregular PCM that prematurely fractures. This phenotype was rescued by elimination of microtubule-dependent pulling forces (via nocodazole) and exacerbated by increasing the magnitude of pulling forces (via *csnk-1(RNAi)*). We conclude that these mutations create structural deficiencies in the PCM scaffold that make it susceptible to fracture under mechanical stress. Our results also demonstrate that mechanical properties must be considered when assaying PCM assembly phenotypes.

The behavior of SPD-5 is typical of associative polymers with a hierarchy of interactions (Choi et al., 2020a; Choi et al., 2020b). In this interpretation, SPD-5’s coiled-coil domains act as “stickers”, while the intervening linker sequences could act as “spacers”(Figure 5). The stickers primarily mediate inter-protein contacts. Spacers engage in limited interactions and likely create intra-protein flexibility, allowing the stickers to sample different configurations. The serial linkage of coiled-coil domains in these proteins creates multivalency, which favors self-assembly through avidity (Li et al., 2012). Association of the coiled-coil stickers yields irregular networks, and certain stickers are more important for assembly than others. But this alone is not sufficient to form spherical PCM seen in cells. Macromolecular crowders, like PEG, can constrain the networks into spherical “condensates” that more closely mimic PCM morphology *in vivo* (Woodruff et al., 2017). Recent theories suggest that crowding agents induce protein phase separation by altering linker solubility (Harmon et al., 2017). Whether this principle applies *in vivo* remains to be tested. SPD-5 interacts directly with several client proteins (e.g., SPD-2, PLK-1, RSA-2 and others)(Holzer et al., 2022; Woodruff et al., 2017; Woodruff et al., 2015; Wueseke et al., 2014), which could influence PCM scaffold assembly and morphology in complex ways (Ruff et al., 2021). Future studies will test the importance of the linker regions and identify how crowding and the many client proteins cooperate to influence PCM assembly, strength, and morphology.

Our data also show that SPD-5 residues interact with various partners during scaffold assembly. This property is not expected for regularly oriented and registered coiled coils; for example, zero-length XL-MS analysis of laminin revealed only single-partner and occasionally double-partner crosslinks (Armony et al., 2016). These data suggest that the assembled SPD-5 scaffold should not exhibit long-range order, consistent with previous cryo-EM analysis (Woodruff et al., 2017; Woodruff et al., 2015). These features could explain why SPD-5 forms micron-scale, super-stoichiometric assemblies rather than stoichiometric complexes typical of coiled-coil proteins (Truebestein and Leonard, 2016).

At face value, our multivalent polymer model seems to be at odds with observations of *spd-5(or213)* embryos, which express SPD-5 harboring a single mutation (R592K) that prevents PCM assembly and causes lethality at 25°C (Hamill et al., 2002). Why should mutation of any single domain dramatically affect assembly of a multivalent polymer? In our companion paper, we address this issue and show that R592 lies within an alpha helical hairpin essential for PCM scaffold strength (Rios et al., 2023). PCM assembly in *spd-5(or213)* embryos is largely rescued by removal of microtubule-mediated forces. Thus, external forces sensitize the system, such that small perturbations cause fracture and disassembly of the SPD-5 scaffold under stress. Similar cases are seen in engineering, where failure of structures can be attributed to local discontinuities (Oliver et al., 2004).

In conclusion, we propose that multivalent coiled-coil interactions between SPD-5 molecules undergird PCM scaffold assembly and impart sufficient strength to resist microtubule-mediated forces in *C. elegans* embryos (Figure 5). PLK-1 potentiates SPD-5 multimerization by relieving non-specific interactions and creating new, perhaps higher affinity interactions between coiled-coil domains. Like SPD-5, PCM scaffold proteins in other species (e.g., Centrosomin, CDK5RAP2, PCNT) contain numerous dispersed coiled-coil domains and PLK-1 phosphorylation sites. It is possible that PCM scaffold assembly through multivalent coiled-coils domains is conserved. More broadly, our work reveals the importance of emergent material properties for organelle function and highlights serial coiled-coil motifs as drivers of micron-scale, super-stoichiometric assemblies in cells. These principles could provide a framework to understand the formation of other supramolecular assemblies rich in coiled-coil proteins, including microtubule-organizing centers, RNP granules, and endocytic initiation sites (Ford and Fioriti, 2020; Kozak and Kaksonen, 2022; Sallee and Feldman, 2021).

## DATA AVAILABILITY

Further requests and information for resources and reagents should be directed to and will be fulfilled by the Lead Contact, Jeffrey Woodruff. Raw mass spectrometry files can be found on the MassIVE database:

Accession number MSV000090165 http://massive.ucsd.edu/ProteoSAFe/status.jsp?task=b36ed704fe8f4ed9b5100da390fe168a

Accession number MSV000090165 http://massive.ucsd.edu/ProteoSAFe/status.jsp?task=3e56f634ef304ce4b0a11366b66bbc15

## Supporting information

Supplemental Data Set 1

Supplemental Data Set 2

Supplemental Data Set 3

## ACKNOWLEDGEMENTS

We would like to thank Anthony Hyman, Karen Oegema, and Jessica Feldman for providing strains and antibodies; Andrea Zinke, Andrey Pozniakovsky, and Susanne Ernst for help with transgenic worm construction; Barbara Borgonovo for help with CD spectroscopy; the Protein Expression Facility at MPI-Dresden for help with expressing proteins; the Proteomics Core Facility at UT Southwestern for mass spectrometry; Jesse Bucksot and Sofia Bali for help with developing MATLAB analysis scripts.

## AUTHOR CONTRIBUTIONS

M.U.R. performed and analyzed *in vivo* experiments involving *C. elegans* embryos, purified dephosphorylated SPD-5, and performed and analyzed XL-MS experiments of phosphorylated SPD-5. M.A.B. purified SPD-5, performed SPD-5 XL-MS experiments, and quantified cross-linking results. B.F-G. purified SPD-5 protein fragments and performed circular dichroism experiments. B.D.R. and L.A.J. analyzed cross-linking data. N.F. performed baculovirus-mediated protein expression and immunofluorescence. K.Y. analyzed mutant embryos expressing RFP::SPD-5 and RFP::SPD-5(Δ734-918). J.B.W. performed *in vitro* assays, optimized XL-MS protocols, and analyzed data. J.B.W. wrote the manuscript.

## MATERIALS AND METHODS

### Experimental model and subject details

For expression of recombinant proteins (listed in Table S1) we used SF9-ESF *S. frugiperda* insect cells grown at 27**°**C in ESF 921 Insect Cell Culture Medium (Expression Systems) supplemented with Fetal Bovine Serum (2% final concentration). *C. elegans* worm strains were grown on nematode growth media (NGM) plates at 16-23**°**C, following standard protocols (www.wormbook.org). Worm strains used in this study are listed in Table S2 and created using MosSCI(Frokjaer-Jensen et al., 2008) or CRISPR(Paix et al., 2015).

### Prediction of coiled-coil motifs and secondary structure

Coiled-coil motifs in SPD-5 were predicted using MARCOIL at 50% threshold, an MTK matrix, and high transition probability (https://toolkit.tuebingen.mpg.de/tools/marcoil). Secondary structure was predicted using Alphafold (Jumper et al., 2021; Varadi et al., 2022).

### Protein purification

All expression plasmids are listed in Table S1. Full-length SPD-5 and PLK-1 constructs were expressed and purified as previously described (Woodruff and Hyman, 2015; Woodruff et al., 2015). For newly made proteins, genes encoding SPD-5 fragments were amplified from the full-length *spd-5* coding sequence using PCR, then inserted into a baculoviral expression plasmid (pOCC27 or pOCC28) using standard restriction cloning. Baculoviruses were generated using the FlexiBAC system (Lemaitre et al., 2019) in SF9 cells. Protein was harvested 72 hr post infection during the P3 production phase. Cells were collected, washed, and resuspended in harvest buffer (25 mM HEPES, pH 7.4, 150 mM NaCl). All subsequent steps were performed at 4**°**C. Cell pellets were resuspended in Buffer B (50 mM Tris, pH 7.4, 30mM imidazole, 500 mM KCl, 0.5 mM DTT, 1% glycerol, 0.1% CHAPS) + protease inhibitors, then lysed using a dounce homogenizer. Proteins were bound to Ni-NTA (Qiagen), washed with 10 column volumes of Buffer A, eluted with 250 mM imidazole. The eluate was then bound to amylose resin (NEB), washed with 5 column volumes of Buffer C (25 mM HEPES, pH 7.4, 500 mM NaCl, 0.5 mM DTT, 1% glycerol, 0.1% CHAPS). Protein was eluted by adding PreScission protease, incubating overnight, and then passed over Glutathione Sepharose 4B (Sigma) to remove the Precission protease. Eluted protein was then concentrated using 3K-50K MWCO Amicon concentrators (Millipore). All proteins were aliquoted in PCR tubes, flash-frozen in liquid nitrogen, and stored at -80**°**C. Protein concentration was determined by measuring absorbance at 280 nm using a NanoDrop ND-1000 spectrophotometer (Thermo Scientific).

### Cross-linking mass spectrometry of SPD-5

To achieve reliable results, we found it was necessary to start with completely dephosphorylated SPD-5. Purification of dephosphorylated SPD-5 was achieved through modification of our standard protocol. In short, insect cell lysates were passed through 4mL of Ni-NTA beads twice, washed five times with buffer 1 (25mM HEPES, 500mM NaCl, 30mM imidazole, 1% glycerol, 0.1% CHAPS, pH7.4), then twice with buffer 2 (150mM KCl, 25mM HEPES, pH7.4) at 4°C. Ni-NTA-bound SPD-5 was incubated for 1 hr at room temperature in dephosphorylation buffer (1X PMP buffer (NEB) + 1mM MnCl_2_, + 40,000 U of lambda phosphatase (400,000 units/mL, NEB). Beads were then washed twice with buffer 1 at 4°C. Dephosphorylated SPD-5 was eluted from the Ni-NTA beads using 15mL of Buffer 3 (25mM HEPES, 500mM NaCl, 250mM imidazole, 1% glycerol, 0.1% CHAPS, pH7.4). SPD-5 was then bound to 500 μL MBP-trap beads (Chromotek) and the column was washed 3X with buffer 4 (25mM HEPES, 500mM NaCl, 1% glycerol, 0.1% CHAPS, pH7.4). Dephosphorylated SPD-5 was eluted from the MBP-trap beads by overnight incubation in buffer 4 + 100μL of PreScission protease (Acro Biosystems; 1mg/mL) at 4°C. Eluted protein was further purified and stored as described above. Dephosphorylation of SPD-5 was confirmed by PTM identification using mass spectrometry showing effective removal of 99.8-100% of phosphates.

Crosslinking reactions were prepared at room temperature with 1μM dephosphorylated SPD-5, 1μM PLK-1 (KD/CA), 0.2 mM ATP, 10 mM MgCl_2_, 150mM KCl, 25mM HEPES, pH7.4 and 0.5mM DTT. Samples were incubated for 2 hr at room temperature followed by addition of 8 mM DMTMM for 45 min at room temperature (shaking at 300 rpm). To quench the reaction, we added 50 mM ammonium bicarbonate for 15 min at room temperature (shaking at 300rpm). Samples without cross-linker were used for mass spectrometry PTM analysis to identify phosphorylated sites (Supplementary Data Set 2).

Samples were run on an SDS-PAGE gel to separate cross-linked species. Bands corresponding to monomeric or multimeric protein were excised from the gel, then digested overnight with trypsin (Pierce), reduced with DTT, and alkylated with iodoacetamide (Sigma). Samples were cleaned using solid-phase extraction with an Oasis HLB plate (Waters), then injected into an Orbitrap Fusion Lumos mass spectrometer coupled to an Ultimate 3000 RSLC-Nano liquid chromatography system. Peptides were separated using a 75 μm i.d., 75-cm long EasySpray column (Thermo) and eluted with a gradient at a flow rate of 250 nL/min from 0-5% buffer B over 1 min, 5%-40% B over 60 minutes, 40%-99% over 25 minutes, and held at 99% B for 5 minutes before returning to 0% B for column equilibration. Buffer A contained 2% (v/v) ACN and 0.1% formic acid in water, and buffer B contained 80% (v/v) ACN, 10% (v/v) trifluoroethanol, and 0.1% formic acid in water. The mass spectrometer operated in positive ion mode with a source voltage of 1.5-2.4 kV and an ion transfer tube temperature of 275°C. MS scans were acquired at 120,000 resolution in the Orbitrap and up to 10 MS/MS spectra were obtained in the ion trap for each full spectrum acquired using collision-induced dissociation (CID) for ions with charges 3-7. Dynamic exclusion was set for 25 s after an ion was selected for fragmentation.

For data analysis, each Thermo.raw file was converted to .mzXML format for analysis using an in-house installation of xQuest (Leitner et al., 2014). Score thresholds were set through xProphet(Leitner et al., 2014), which uses a target/decoy model. The search parameters were set as follows. For zero-length cross-link search with DMTMM: maximum number of missed cleavages = 2, peptide length = 5–50 residues, fixed modifications carbamidomethyl-Cys (mass shift = 57.02146 Da), mass shift of cross-linker = −18.010595 Da, no monolink mass specified, MS^1^ tolerance = 15 ppm, and MS^2^ tolerance = 0.2 Da for common ions and 0.3 Da for cross-link ions; search in enumeration mode. For grouping heavy and light scans (DSS crosslinks only): precursor mass difference for isotope-labeled succinimides = 12.07573 Da for DSS-h_12_/d_12_; maximum retention time difference for light/heavy pairs = 2.5 min. Maximum number of missed cleavages (excluding the cross-linking site) = 2, peptide length = 5–50 aa, fixed modifications = carbamidomethyl-Cys (mass shift = 57.021460 Da), mass shift of the light crosslinker = 138.068080 Da, mass shift of mono-links = 156.078644 and 155.096428 Da, MS^1^ tolerance = 10 ppm, MS^2^ tolerance = 0.2 Da for common ions and 0.3 Da for cross-link ions, search in enumeration mode. The false discovery rates (FDR) for all experiments were <20% at the link level (see Supplemental Data Sets 3 for further information about cross-linked peptide pairs). Samples were also re-analyzed accounting for mass shifts due to the presence of phospho-serines and phospho-threonines. A cumulative list of cross-linked pairs can be found in Supplementary Data Set 1.

### RNAi treatment

RNAi was done by feeding. The *spd-5* feeding clone targets a region that is reencoded in our MosSCI transgenes (Woodruff et al., 2015). Bacteria were seeded onto nematode growth media (NGM) supplemented with 1 mM isopropyl b-D-1-thiogalactopyranoside (IPTG) and 100 µg mL^-1^ ampicillin. L4 hermaphrodites were grown on feeding plates at 23°C for 24 hours.

### Western Blotting

60 adult worms were picked and transferred to blank plates for 20 min to remove bacteria off their bodies. Worms were then moved to PCR tubes containing 10µL of mili-Q water to which 10µL of SDS loading buffer was added. Samples were separated by SDS-PAGE. Protein from each gel was transferred to a nitrocellulose membrane using a Trans-blot turbo transfer for high molecular weight proteins (10 minutes). Membranes were incubated in blocking buffer consisting of 1X TBS-T + 3% Blotting-Grade Blocker (BioRad) shaking at room temperature for 1 hr. Membranes were then washed three times with fresh 1X TBS-T and incubated shaking with primary antibodies over night at 4°C. Primary antibody was washed three times with fresh 1X TBS-T and incubated shaking with secondary antibodies at room temperature for one hour. Each membrane was then incubated in ECL reagent (Thermo Scientific SuperSignal West Femto) for 5 minutes and imaged with a ChemiDoc Touch Imaging System. <u>Primary Antibodies:</u> Mouse anti-alpha tubulin (1:1,000; Cell Signaling Technologies, Product # 3873S Lot #15); Goat anti-enhanced GFP (1:5,000; Dresden PEP facility, 15mg/mL), Rabbit anti-SPD-5 (1:1,000, clone 758, Dresden Antibody Facility). <u>Secondary Antibodies (1:50,000 for all):</u> HRP conjugated Goat anti-Rabbit IgG (1mg/mL) (Invitrogen, # 65-6120, Lot # J276300); HRP conjugated Goat anti-Mouse (1.5 mg/mL) (Invitrogen # 62-6520, Lot # WA312227); HRP conjugated Donkey anti-Goat (1mg/mL) (Invitrogen #A15999, Lot # 58-155-072318).

### Embryo viability assay

10 L4 worms picked and transferred to OP50 plates. 24 hours later, individual adult worms were transferred to individual plates (10 worms per strain) and allowed to lay eggs for 6 hr. Adult worms were then removed from the mating plates and eggs were manually counted. Hatched worms were then counted 24 hr later. Plate viability is reported as number of worms hatched divided by number of eggs laid. Strain viability is reported as the average viability of 10 plates.

### Generation of C. elegans embryos expressing spd-5 transgenes

*C. elegans* worm strains used in this study were created using MosSCI(Frokjaer-Jensen et al., 2008) and based on constructs made previously(Woodruff et al., 2015). Briefly, genomic sequences representing *spd-5* fragments were amplified from pOD1021(MosII)*Pspd2::spd-5(1-1198)::3’UTR* and inserted into the pCFJ151-based parent plasmid (pOD1021(MosII)) lacking the full-length insert. The plasmids were purified using a NucleoBond Xtra Midi Prep Kit (Macherey Nagel), combined with co-injection plasmids, and injected into strain EG6699 (tt5605, Chr II). After one week, worms were heat-shocked for 3 hr at 35℃ to kill worms maintaining extrachromosomal arrays. Moving worms without fluorescent co-injection markers were selected as candidates. Sequencing was used to confirm transgene integration.

### Generation of CRISPR-modified C. elegans mutants

*C. elegans* worms expressing tagRFP::SPD-5 at the endogenous spd-5 locus (gift of J. Feldman) were modified by CRISPR-Cas9 to delete the 555 bp sequence encoding a.a. 734-918 (*PHX5737 spd-5(syb5737))*. Modified worms were generated by SunyBiotech.

### Microscopy of *C. elegans* embryos

Embryos from adult worms were dissected on a 22 × 50 mm coverslip (Coring Catalog # 2975-225) containing 20µL of egg salts buffer (ESB) with 15-µm polystyrene beads (Sigma) using two 22-gauge needles. Samples were then mounted onto plain 25 x 75 x 1 mm microscope slides (Fisher # 12-544-4). Time-lapse images were acquired with an inverted Nikon Eclipse Ti2-E microscope with a Yokogawa confocal scanner unit (CSU-W1), piezo Z stage, and an iXon Ultra 888 EMCCD camera (Andor), controlled by Nikon Elements software. We used a 60× 1.2-NA Plan Apochromat water-immersion objective to acquire 41 × 0.5-µm Z-stacks with 488 nm excitation (15 % laser power), 100 ms exposure time, 1 min intervals, using 2 × 2 binning followed by DIC imaging (92.3% iris intensity) using 1 x 1 binning.

### *In vitro* condensate assays

To assess SPD-5 self-assembly, GFP-labeled SPD-5 was incubated in condensate assembly buffer (25 mM HEPES, 150mM KCl, 0.5 mM DTT, 3-9% (w/v) PEG-3350). To assess recruitment, 1000 nM SPD-5::RFP was incubated in condensate assembly buffer for 2 min, then 10 nM SPD-5::GFP (final concentration) was added. All condensates were transferred to a pre-cleaned glass bottom imaging plate (Corning ref#4580; 96 well) and settled for 5 min before imaging with a Nikon Eclipse Ti-2E spinning disk confocal microscope (described in the live-cell imaging section) and either a 40X 1.25 NA silicone or 100X 1.35 NA silicone immersion objective.

### Circular dichroism

0.3 mg/ml full-length SPD-5 or SPD-5 fragments were incubated in buffer (20 mM K-Phosphate, pH 7.4, 0.01% CHAPS, 0.05 mM DTT, 100 mM NaCl) for 5 min at room temperature before being loaded into a 0.5 mm quartz cuvette (Hellma). Samples were analyzed using a Chirascan CD Spectrometer (Applied Photophysics).

### Embryo immunofluorescence

Embryos were collected from adult *C. elegans* fed on *spd-5(RNAi)* plates for 24 hr. Embryos were mounted in M9 on Superfrost Plus slides (Fisher) and frozen in liquid nitrogen, fixed with methanol at -20°C for 10 min, then washed twice with TBS-T for 5 min. Samples were blocked in TBST +3% BSA for 30 min at room temperature, then incubated 1 hr with blocking buffer + primary antibodies (1:1000 rabbit anti-SAS-4(Kirkham et al., 2003), 1:1000 rat anti-alpha tubulin-alexa647(Invitrogen), and 1:5000 rabbit anti-SPD-5(lot 785)). Slides were washed 3 times for 5 min with TBST, incubated for 1 hr in blocking buffer + secondary antibodies (1:1000 donkey anti-rabbit alexa555 (Invitrogen); goat anti-rabbit alexa488(Invitrogen)), then washed 3 times for 5 min with TBS-T. Cells were mounted in Vectashield mounting medium with DAPI (Vector laboratories) and imaged by fluorescence confocal microscopy (100X 1.35 NA silicone objective).

### Image quantification and statistical analyses

Images were analyzed using semi-automated, threshold-based particle analysis in FIJI. Data were plotted and statistical tests were performed using GraphPad prism. The sample size, measurement type, error type, and statistical test are described in the Figure legends where appropriate.

## Supplementary Tables

**TABLE S1.** Constructs for protein expression.

| protein | sequence | plasmid name | parent plasmid (pOCC #) | N-term tag | C-term tag |
| --- | --- | --- | --- | --- | --- |
| SPD-5 | Full-length | JWV2 | pOCC28 | MBP-PreScission | PreScission-6xHis |
| SPD-5::GFP | Full-length | JWV1 | pOCC27 | MBP-PreScission | eGFP-PreScission-6xHis |
| SPD-5::RFP | Full-length | JWV3 | pOCC25 | MBP-PreScission | tagRFP-PreScission-6xHis |
| SPD-5 (F20) | aa 1-386 | JWV20 | pOCC27 | MBP-PreScission | eGFP-PreScission-6xHis |
| SPD-5 (F21) | aa 1-566 | JWV21 | pOCC27 | MBP-PreScission | eGFP-PreScission-6xHis |
| SPD-5 (F22) | aa 281-386 | JWV22 | pOCC27 | MBP-PreScission | eGFP-PreScission-6xHis |
| SPD-5 (F23) | aa 386-566 | JWV23 | pOCC27 | MBP-PreScission | eGFP-PreScission-6xHis |
| SPD-5 (F24) | aa 566-1198 | JWV24 | pOCC27 | MBP-PreScission | eGFP-PreScission-6xHis |
| SPD-5 (F25) | aa 730-990 | JWV25 | pOCC27 | MBP-PreScission | eGFP-PreScission-6xHis |
| SPD-5 (F26) | aa 730-1198 | JWV26 | pOCC27 | MBP-PreScission | eGFP-PreScission-6xHis |
| SPD-5 (F27) | aa 990-1198 | JWV27 | pOCC27 | MBP-PreScission | eGFP-PreScission-6xHis |
| SPD-5 (ΔCC-LONG) | a.a. 734-918 deleted | JWV118 | pOCC27 | MBP-PreScission | eGFP-PreScission-6xHis |
| SPD-5 (F20) | aa 1-386 |  | pOCC28 | MBP-PreScission | PreScission-6xHis |
| SPD-5 (F21) | aa 1-566 |  | pOCC28 | MBP-PreScission | PreScission-6xHis |
| SPD-5 (F22) | aa 281-386 |  | pOCC28 | MBP-PreScission | PreScission-6xHis |
| SPD-5<br>(F23) | aa 386-566 |  | pOCC28 | MBP-PreScission | PreScission-6xHis |
| SPD-5<br>(F24) | aa 566-1198 |  | pOCC28 | MBP-PreScission | PreScission-6xHis |
| SPD-5<br>(F25) | aa 730-990 |  | pOCC28 | MBP-PreScission | PreScission-6xHis |
| SPD-5<br>(F26) | aa 730-1198 |  | pOCC28 | MBP-PreScission | PreScission-6xHis |
| SPD-5<br>(F27) | aa 990-1198 |  | pOCC28 | MBP-PreScission | PreScission-6xHis |
| PLK-1(CA) | T194D | JWV11 | pOCC7 | 6xHis-PreScission |  |
| PLK-1(KD) | K67M | JWV12 | pOCC7 | 6xHis-PreScission |  |

**TABLE S2.**
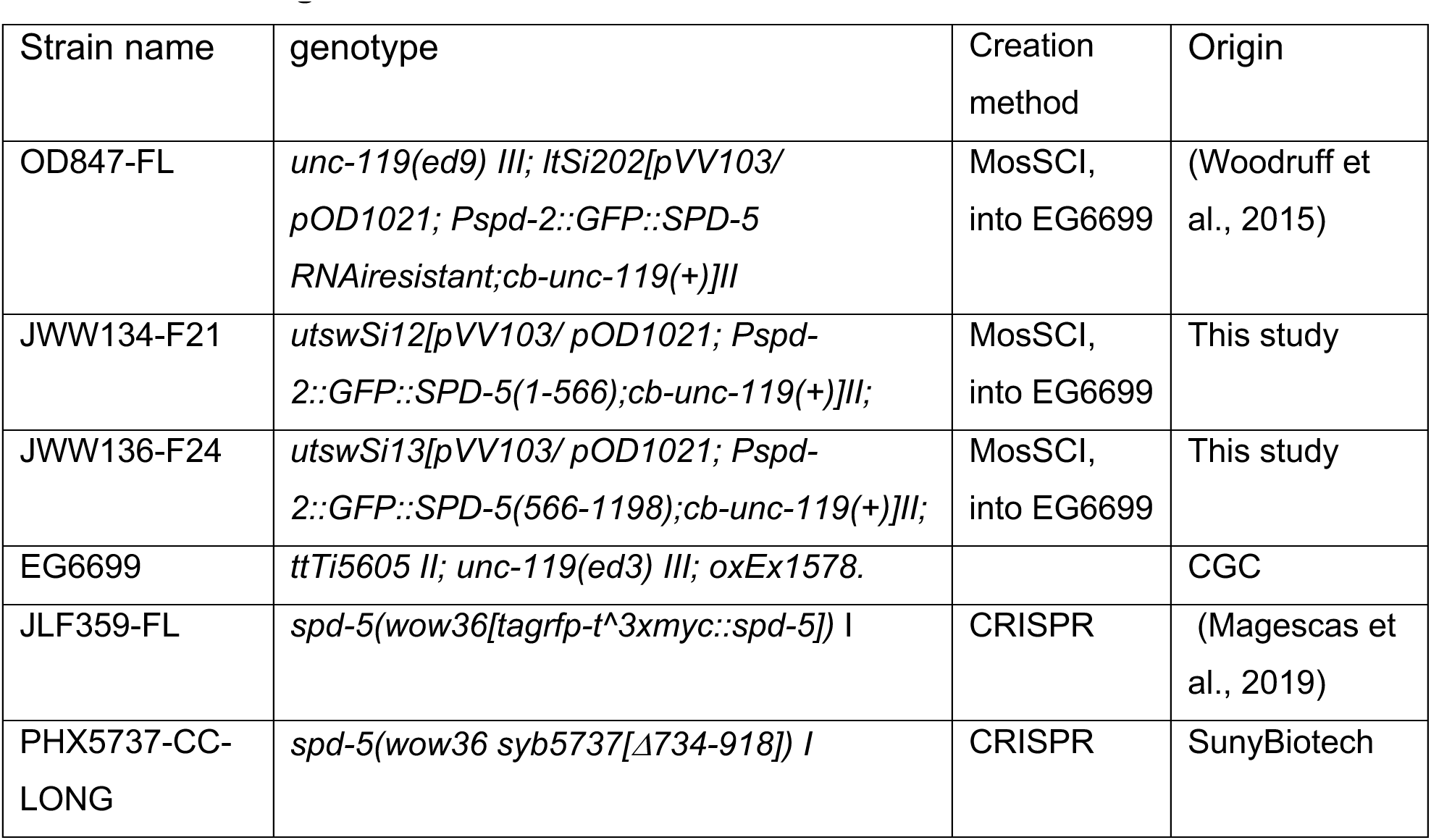
C. elegans strains.

## Supplementary Data

**Supplementary data set 1. Cumulative list of interacting pairs identified by XL-MS.** See attached .xls file.

**Supplementary data set 2. Identified phosphorylated sites on SPD-5.** See attached .xls file.

**Supplementary data set 3. XL-MS peptide analysis of SPD-5 with PLK-1 (CA) and PLK-1 (KD).** See attached .xls file.

## SUPPLEMENTAL FIGURE LEGENDS

**Figure S1.**
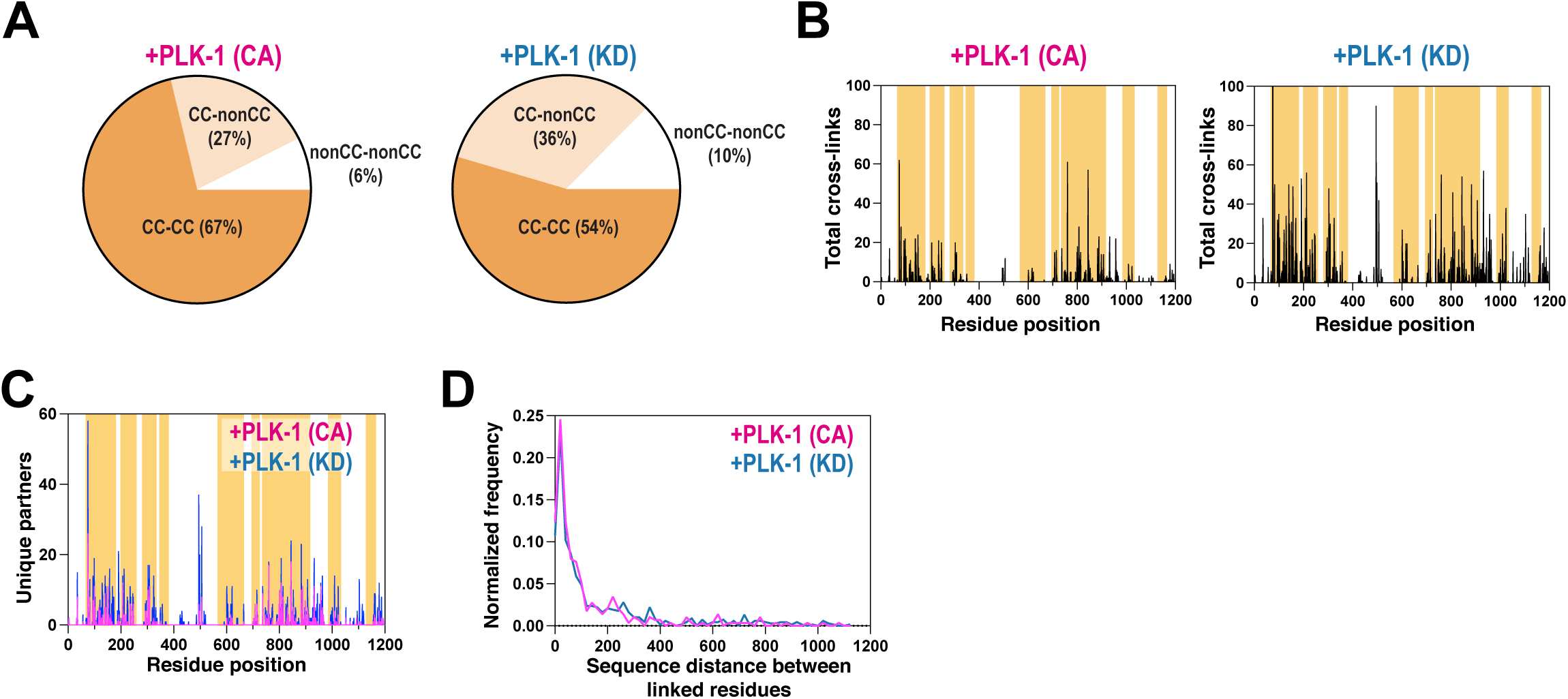
Analysis of cross-linked residues in phosphorylated and unphosphorylated SPD-5. A. Percentage of cross-linked pairs involving predicted coiled-coil domains (CC) or linker domains from multimeric SPD-5 samples incubated with constitutively active PLK-1 (CA) or kinase dead PLK-1 (KD) (n = 6 replicates for each condition). B. Map showing total number of times a residue was identified in a cross-linked pair. Predicted coiled-coil domains are indicated in orange. C. For each residue in SPD-5, the number of unique cross-linked partner residues was calculated. D. Histogram of sequence distances (in residues) between cross-linked pairs. Data were grouped into bins of 20 residues.

**Figure S2.**
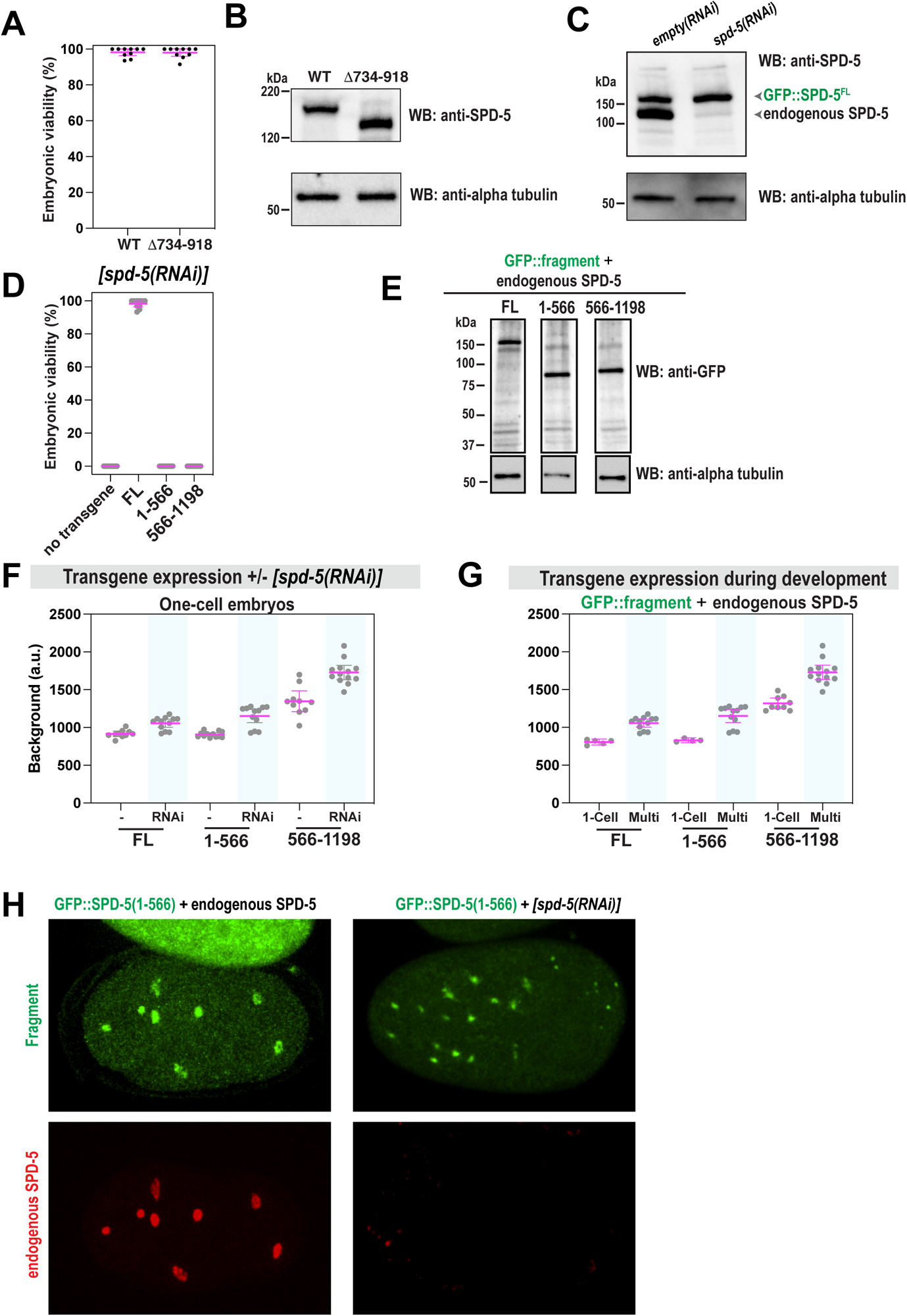
Control experiments for SPD-5 mutant analysis. A. Viability of offspring after mothers were grown on normal OP50 bacteria. Embryos expressed either RFP::SPD-5(WT) or RFP::SPD-5(Δ734-918). Mean +/- 96% C.I.; n = 10 worms per condition, 23-44 embryos each. B. Western blot showing expression of RFP::SPD-5(WT) or RFP::SPD-5(Δ734-918). Alpha tubulin was used as a loading control. C. Western blot showing depletion of endogenous SPD-5, but not transgenic GFP::SPD-5, following RNAi. Alpha tubulin was used as a loading control. D. Viability of offspring after mothers were fed for 24 hr on *spd-5(RNAi)* plates. No transgene (N2) compared with MosSCI worms expressing transgenic *gfp::spd-5*. Mean +/- 95% C.I.; n = 10 worms per condition, 25-40 embryos each. E. Western blot showing expression of *gfp::spd-5* transgenes. Alpha tubulin was used as a loading control. F. Quantification of cytoplasmic fluorescence of transgenic GFP::SPD-5 proteins in one-cell embryos with and without endogenous SPD-5. Mean +/- 95% C.I.; n = 7-13 embryos. G. Quantification of cytoplasmic fluorescence of transgenic GFP::SPD-5 proteins in in One-cell stage vs. multi-cell stage embryos (8-cell or greater). Mean +/- 95% C.I.; n = 5-12 embryos). H. Immunofluorescence of transgenic embryos grown on control or *spd-5(RNAi)* feeding plates. GFP signal was preserved by light fixation. Endogenous SPD-5 was detected by an antibody that targets its C-terminus.

**Figure S3.**
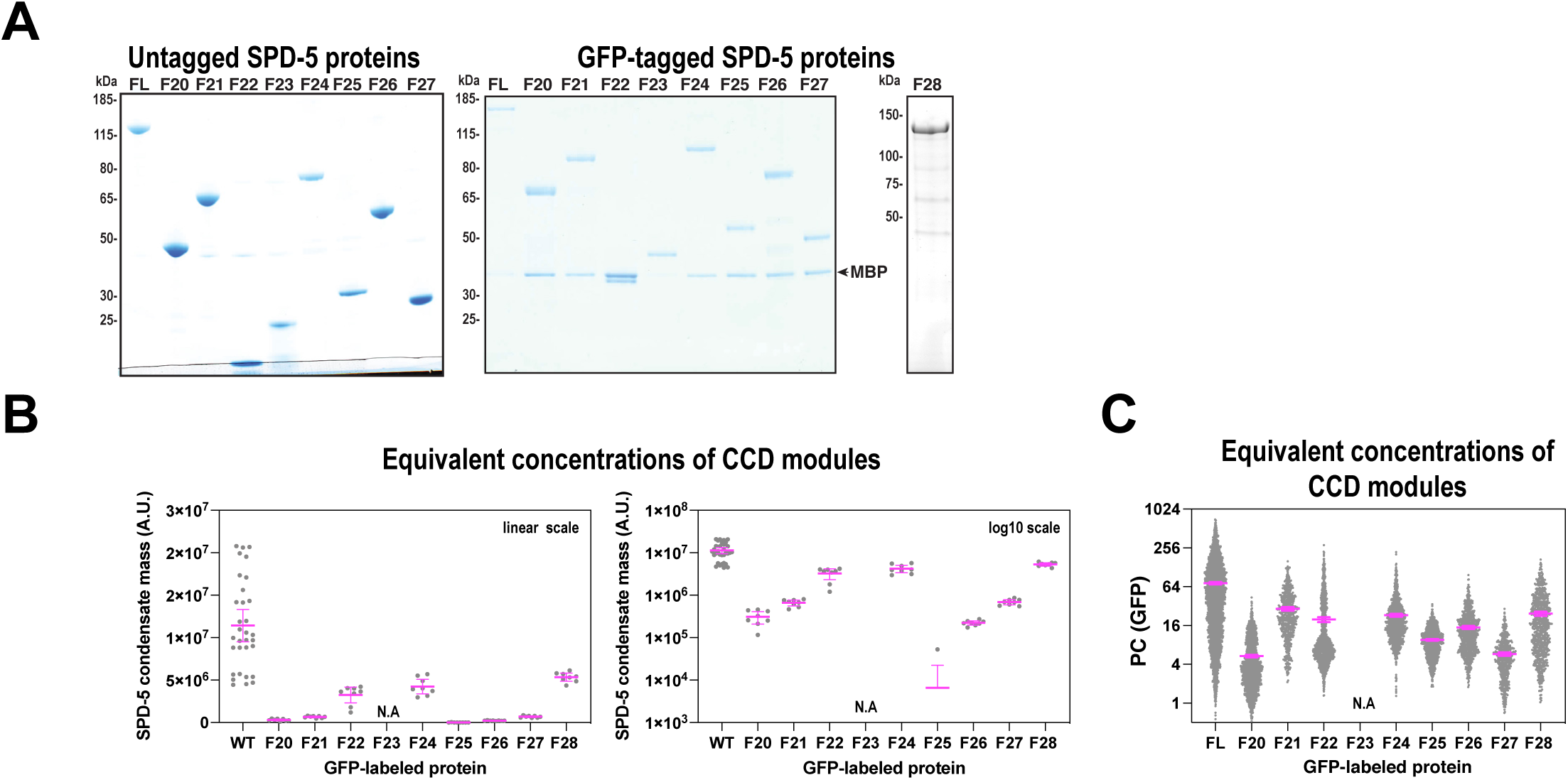
Analysis of SPD-5 assembly and recruitment *in vitro*. A. Coomassie-stained SDS-PAGE gels of SPD-5 proteins used in this study. B. *in vitro* SPD-5 assembly using equivalent molar concentrations of coiled-coil mass (500 nM (FL), 1140 nM (F20), 1140 nM (F21), 3315 nM (F22), 890 nM (F24), 1675 nM (F25), 1120 nM (F26), 3390 nM (F27), 710 nM (F28). Mean +/- 95% C.I.; n = 11-22 images. C. *in vitro* SPD-5 recruitment using equivalent molar concentrations of coiled-coil mass (1000 nM SPD-5::RFP; for GFP proteins: 10 nM (FL), 23 nM (F20), 23 nM (F21), 66 nM (F22), 18 nM (F24), 36 nM (F25), 22 nM (F26), 68 nM (F27), 14 nM (F28). Mean +/- 95% C.I. n = 532-4702 condensates.

